# A sparse observation model to quantify species interactions in time and space

**DOI:** 10.1101/815027

**Authors:** Sadoune Ait Kaci Azzou, Liam Singer, Thierry Aebischer, Madleina Caduff, Beat Wolf, Daniel Wegmann

## Abstract

Camera traps and acoustic recording devices are essential tools to quantify the distribution, abundance and behavior of mobile species. Varying detection probabilities among device locations must be accounted for when analyzing such data, which is generally done using occupancy models. We introduce a Bayesian Time-dependent Observation Model for Camera Trap data (`Tomcat`), suited to estimate relative event densities in space and time. `Tomcat` allows to learn about the environmental requirements and daily activity patterns of species while accounting for imperfect detection. It further implements a sparse model that deals well will a large number of potentially highly correlated environmental variables. By integrating both spatial and temporal information, we extend the notation of overlap coefficient between species to time and space to study niche partitioning. We illustrate the power of `Tomcat` through an application to camera trap data of eight sympatrically occurring duiker *Cephalophinae* species in the savanna - rainforest ecotone in the Central African Republic and show that most species pairs show little overlap. Exceptions are those for which one species is very rare, likely as a result of direct competition.

## 1. Introduction

Thanks to their automated and non-intrusive nature of observation, camera traps, acoustic recorders and other devices that allow for continuous recording of animal observations have become an essential part of many wildlife monitoring efforts, especially those that aim at quantifying the distribution, abundance and behavior of mobile species (e.g. O’Brien et al., 2010; Burton et al., 2015; Caravaggi et al., 2017). However, the inference of these biological characteristics is not trivial due to the confounding factor of detection, which may vary greatly among recording locations. Animals are, for instance, more likely to trigger a picture when passing a camera trap in the open savanna than in a dense rainforest. Hence, variation in the rates at which a species is recorded (e.g. the photographic rate) may reflect differences in local abundance, but might just as well reflect differences in the probabilities with which individuals are detected, or more likely a combination of both (see Burton et al., 2015; Sollmann, 2018, for two excellent reviews). Accounting for varying detection rates is thus critical when comparing species densities between locations.

Since local detection rates are generally not known, they have to be inferred jointly with abundance. Commonly used methods to do so are variants of occupancy models that include detection probabilities explicitly (MacKenzie et al., 2002). The basic quantity of interest in these models is whether or not a particular site is occupied by the focal species. While the detection of a species implies that it is present, the absence of a record does not necessarily imply the species is absent. Since the probabilities of detection and occupation are confounded, they can not be inferred for each site individually. Detection probabilities are thus either assumed to be constant across sites or, more commonly, assumed to be a function of environmental covariates governed by hierarchical parameters (MacKenzie et al., 2002).

A problem of occupancy models is the assumption that there exists a well defined patch or site that is either occupied by a species, or not (closure assumption; MacKenzie et al. (2002)). However, the notation of a discrete patch is often difficult, particularly in the case of mobile species that move between camera trap sites, complicating the interpretation of occupied versus empty sites (Efford and Dawson, 2012; Steenweg et al., 2018). In addition, summarizing such data by a simple presence-absence matrix ignores the information about differences in population densities and activities at occupied sites. Occupancy is therefore not necessarily a good surrogate for abundance (Efford and Dawson, 2012; Steenweg et al., 2018; MacKenzie and Royle, 2005), even though it has been advocated for birds (MacKenzie and Nichols, 2004), and identifying an alternative measure has previously been highlighted as a key challenge in wildlife surveys (Burton et al., 2015).

To address this issue, we here introduce `Tomcat`, a Time-dependent Observation Model for Camera Trap data, that extends currently used occupancy models in three important ways:

First, we propose to quantify the rate at which animals pass through a specific location, rather than occupancy. This measure does not easily allow for an absolute quantification of density because it is not possible to distinguish mobility from abundance. But it can be readily compared in space to identify important habitat for a particular species, or in time to identify changes in abundance or activity. It therefore appears more useful than occupancy to monitor changes in species abundances over time, as for many species, changes in population size will be reflected in the rate at which individuals are detected prior to local extinction.

Second, we explicitly model daily activity patterns. Several models have been proposed to estimate such patterns from continuous recording data (Frey et al., 2017), including testing for non-random distributions of observations in predefined time-bins (Bu et al., 2016) and circular kernel density functions (Oliveira-santos et al., 2013; Rowcliffe et al., 2014), with the latter allowing for the quantification of activity overlap between species (Ridout and Linkie, 2009). Jointly inferring activity patterns with the rate at which animals pass a location allows us not only to account for imperfect detection, but also to quantify overlap between species in both time and space, shedding additional light on species interactions.

Third, we explicitly account for the sparsity among environmental coefficients. This is relevant since many environmental covariates are available and it is usually not known which ones best explain the variation in abundance of a species (Kriticos et al., 2014; Title and Bemmels, 2018). Enforcing sparsity on the vector of coefficients avoids the problem of over-fitting in case the number of recording locations is smaller or on the same order as the number of environmental coefficients.

In this article, we begin by describing the proposed model in great detail. We then verify its performance using extensive simulations and illustrate it by inferring spatio-temporal overlap of eight duiker species of the subfamily *Cephalophinae* within the forest-savanna ecotone of Central Africa.

## 2. Material and Methods

### 2.1 A Time-dependent Observation Model

We begin by describing an observation model for continuous recording devices. An illustration of the model is shown in Figure 1.

**Figure 1:**
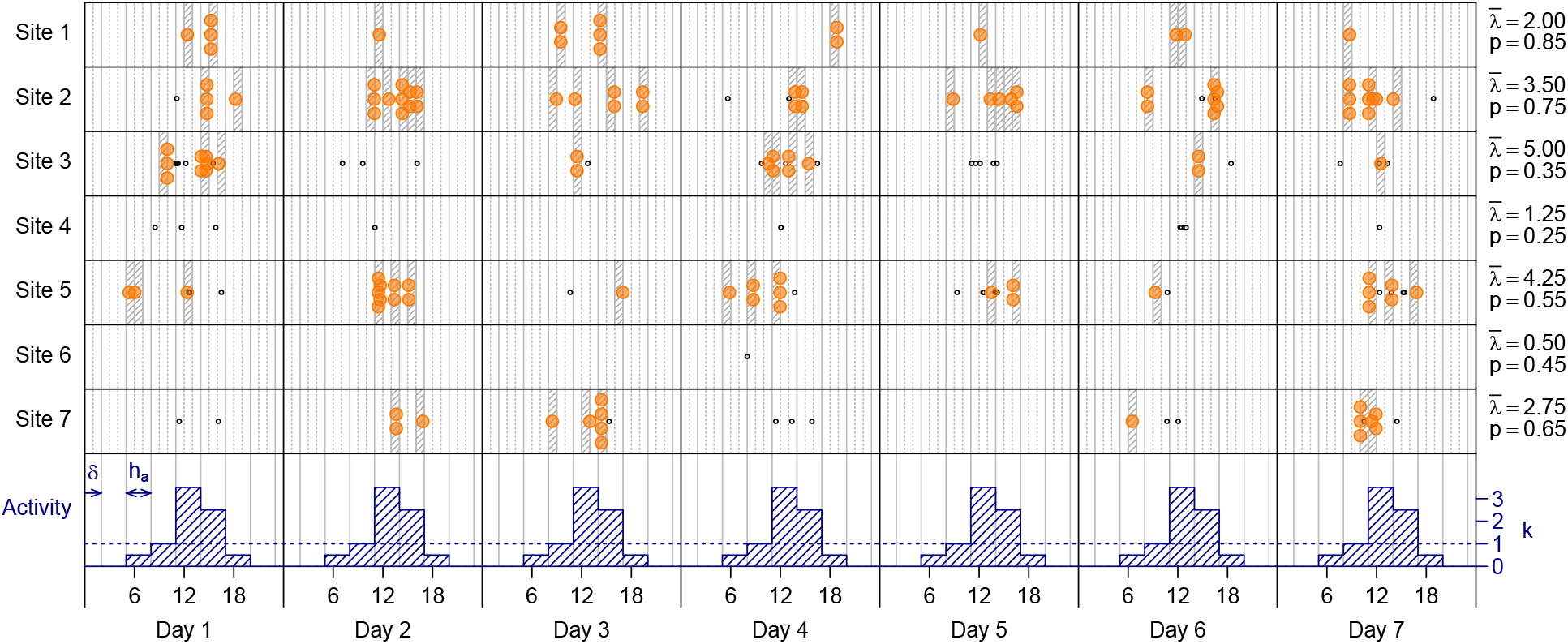
Conceptual plot illustrating the model. Shown are recording devices at seven sites that differ in the rate at which animals pass 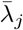 and their detection probabilities *p_j_*. Passing animals may result in multiple records (stacked orange circles) or not be detected (black circles). Boxes illustrate the observation intervals with *h_o_* = 1h and are hatched if at least one animal was recorded within (*W_j_*(·) > 0). The rate at which animals pass is modulated by the daily activity patterns (shown at the bottom) parameterized as a piecewise-constant function 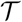 with *h_a_* = 3h, *k* = (0,0.5, 1, 3.5, 2.5, 0.5, 0, 0) and shifted by *δ* = 2h, peaking around mid-day.

Let Λ_*j*_ (*τ*) be the event density at time *τ*: the rate at which a device at location *j* = 1,…, *J* takes observations of a particular species (or guild) at the time of the day *τ* ∈ [0, *T*], *T* = 24h. We assume that this rate is affected by three processes: 1) the average rate 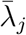 at which individuals pass through location *j*, 2) the daily activity patterns 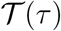 that reflect differences in activity throughout the day, and 3) the probability *p_j_* with which an individual passing through location *j* is recorded (i.e. detected by the device and properly identified downstream):

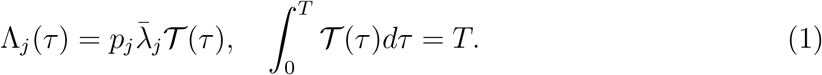

The number of records *W_j_*(*d*,*τ*_1_,*τ*_2_) taken by a device at location *j* within the interval [*τ*_1_,*τ*_2_) on day *d* is then given by the non-homogeneous Poisson process

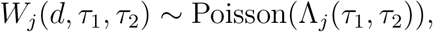

with intensity function

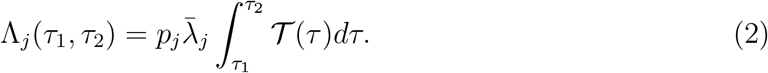

As in all models dealing with imperfect detection, the parameters related to species densities 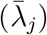 and detection (*p_j_*) are confounded and can not be estimated individually for each location without extra information. However, it is possible to estimate relative densities between locations using a hierarchical model. Following others (e.g. Tobler et al., 2015), we assume that both parameters are functions of covariates (e.g. the environment), and hence only attempt to learn these hierarchical parameters. Here, we use

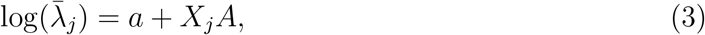

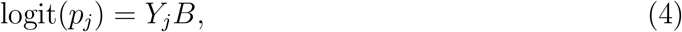

where *X_j_* and *Y_j_* are known (environmental) covariates at location *j* and *a, A* and *B* are species specific coefficients. Note that to avoid non-identifiability issues, we did not include an intercept for pj, and hence we set the average detection probability across locations 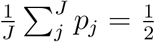 (assuming *X* and *Y* have mean zero). Also, *X* should not contain covariates that are strongly correlated with covariates in *Y* (see below).

#### 2.1.1. Non-independent events

Another issue specific to data from continuous recordings is that not every record is necessarily reflective of an independent observation as the same individual might trigger multiple observations while passing (or feeding, resting, …) in front of a recording device. It is often difficult and certainly laborious to identify such recurrent events. We thus propose to account for non-independent events by dividing the day into *n_o_* intervals of equal length 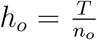 (*o* for observation), and then only consider whether or not at least one recording was taken within each interval [*c*_*m*-1_,*c*_*m*_),*m* = 1,…,*n_o_*, where *c*_0_ = *c_n_o__* = *T* (Figure 1). Specifically, for an interval *m*,

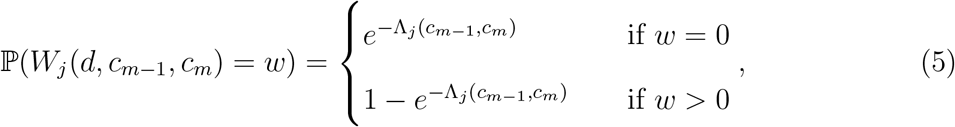

where Λ_*j*_ (*c*_*m*-1_, *c_m_*) is given by (2). We note that the choice of *n_o_* must reflect the activity of the species considered: intervals must be large enough such that it is unlikely that the same individual is recorded in multiple intervals. Too large intervals, however, may impact power as many independent events will end up in the same interval.

#### 2.1.2. Daily activity patterns

Here we assume that 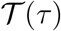 is a piece-wise constant function (or step function) with *n_a_* activity intervals of equal length 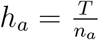 (*a* for activity), i.e. we assume that the activity is constant within a specific interval but independent between intervals (Figure 1). While activity patterns are unlikely strictly piece-wise constant, we chose this function over a combination of periodic functions (e.g. Oliveira-santos et al., 2013) as they fit to complicated, multi-peaked distributions (e.g. crepuscular activity) with fewer parameters.

We denote by *k_i_* the relative activity in interval *i* = 1,…, *n_h_* with *k_i_* = 0 implying no activity and *k_i_* = 1 implying average activity. Since the best way to place the activity intervals within the day (i.e. the tiling) is unknown, we allow for a shift *δ* such that the first interval is [*δ, h_a_* + *δ*) and the last overlaps midnight and becomes [*T − h_a_* + *δ, δ*) (Figure 1). We therefore have

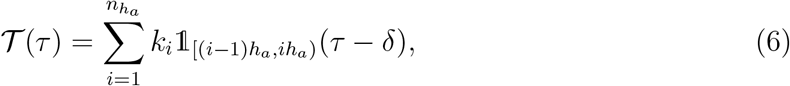

where the indicator function 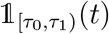 identifies the interval *i* within which *t* falls: it this 1 if *t* ∈ [*τ*_0_, *τ*_1_) and zero otherwise. Note that

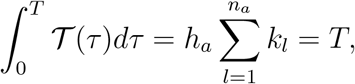

and hence

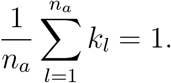

#### 2.1.3. Bayesian inference

We conduct Bayesian inference on the parameter vector ***θ*** = {*a, A, B, δ, **k***}, where ***k*** = {*k*_1_,…, *k_n_a__*}, by numerically evaluating the posterior distribution 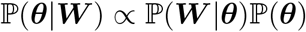, where ***W*** = {*W*_1_,…, *W_J_*} denotes the full data from all locations *j* = 1,…,*J*.

The likelihood 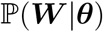 is calculated as

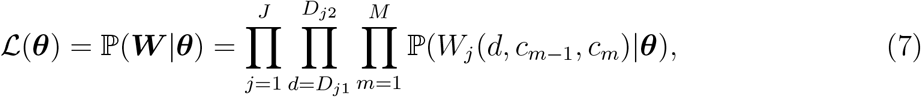

where 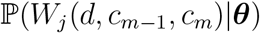 is given by equation (5) and *D*_*j*1_ and *D*_*j*2_ denote the first and last day of recording at location *j*.

##### Prior distribution

Since it is usually not known which covariates *X* and *Y* are informative, nor at which spatial scale they should be evaluated, the potential number of covariates considered may be large. To avoid overfitting, we enforce sparsity on the vectors of coefficients *A* and *B*. Specifically, we introduce the indicators that indicate whether covariate *X_i_* is included in the model (*γ_λi_* = 1), or not (*γ_λi_* = 0). We then assume that *A_i_* follows a normal mixture model such that

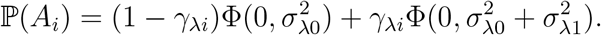

Here, Φ(0,*σ*^2^) denotes the normal density centered at zero with variance *σ*^2^ and the formulation ensures that this variance is larger in the case *γ_λi_* = 1 than in the case *γ_λi_* = 0. We further assume exponential prior distributions *σ*_λ0_ ~ Exp(r_0_) and *σ*_λ1_ ~ Exp(*r*_1_), a Bernoulli prior 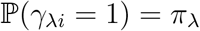, and an analogous prior on *B_i_* with indicators *γ_pi_* and parameters *σ*_*p*0_, *σ*_*p*1_ and *π_p_*. Here we set *r*_0_ = 10^3^ and *r*_1_ = 2.

The parameters *π_λ_* and *π_p_* can only be estimated from data sets with many more sites than environmental covariates. For these cases, we used exponential priors *π_λ_,π_p_* ~ Exp(*r_π_*) and set *r_π_* = 5. If the number of camera trap sites is small or on the same order as the number of environmental covariates, *π_λ_* and *π_p_* should be given *a priori*.

The priors for the remaining parameters were as follows: We chose a normal prior with density 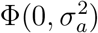 on *a* and set 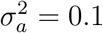. We chose uniform, improper priors on ***k*** and *δ*, namely 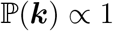 for all vectors of **k** that satisfy 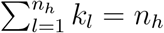, and 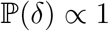 for all 0 ≤ *δ* < *h_a_*. For simplicity, we only consider cases in which *h_a_*, the length of the activity intervals, is a multiple of *h_o_*, the length of the observation intervals, and allow for discrete *δ* ∈ {0, *h_o_*, *2h_o_*,…, *h_a_* − *h_o_*} only.

##### MCMC

We use an MCMC algorithm with Metropolis-Hastings updates to generate samples from the posterior distribution 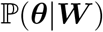, as detailed in the Supporting Information. We run this MCMC for 10,000 iterations after ten successive burnins of 1,000 iterations each. We found these settings to result in proper convergence as assessed using the Gelman-Rubin convergence diagnostics (Gelman and Rubin, 1992) on multiple independent runs (Supplementary Figure S.1).

#### 2.1.4. Prediction

Using a set of *M* posterior samples 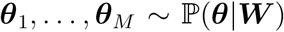, we project event densities to a not-surveyed location 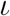 with covariates 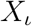 by calculating the mean 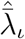 of the posterior 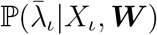 as

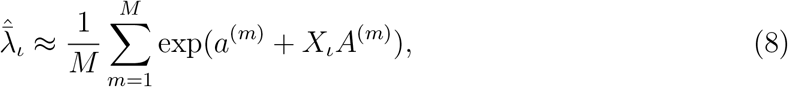

where *a*^(*m*)^ and *A*^(*m*)^ denote the m-th posterior sample of these parameters.

#### 2.1.5. Species overlap in space and time

An important interest in ecology is to compare activity patterns among species and to see how overlapping patterns may relate to their interaction such as competition or predation (e.g. Ridout and Linkie, 2009; Rowcliffe et al., 2014). For that purpose, Ridout and Linkie (2009) introduced overlap coefficients

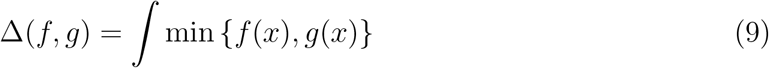

that quantify the overlap between the activity patterns of two species *f* (*x*) and *g*(*x*), respectively, and range from 0 (no overlap) to 1 (identical activity patterns). The overlap Δ(*f,g*) is related to the distance measure *L*_1_ as

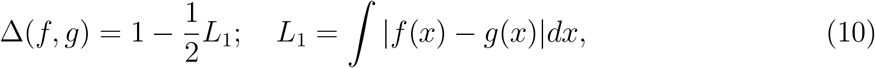

which justifies their visualization of overlap coefficients between *k* species using a Multidimensional Scaling (MDS) by considering 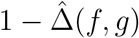 as a measure of dissimilarity.

In practice, the true density functions *f*(*x*) and *g*(*x*) are usually not known. Here we obtain an estimate of Δ numerically from samples *m* = 1,…, *M* of the species-specific posterior distributions obtained with `Tomcat`. We distinguish three types of overlap coefficients: Δ_*T*_ for overlap in time, Δ_*S*_ for overlap in space and Δ_*ST*_ for overlap in time and space. While Δ_*T*_ matches that of Ridout and Linkie (2009), the latter two are extensions made possible by the joint inference of habitat use 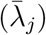 and activity patterns 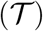 and informative about niche partitioning in space and time.

##### Overlap coefficient Δ_T_

For a large number *n_T_* of equally spaced time values *τ*_1_,*τ*_2_,…, *τ*_*n_T_*_ ∈ (0,*T*), we sample 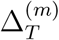 from the posterior distribution 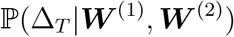 where 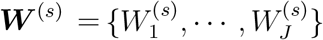 denotes the full data for a species *s* = 1, 2.

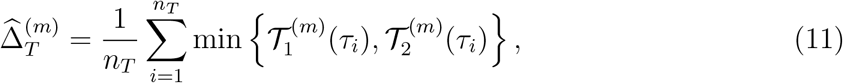

where 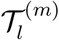 is computed according to equation (6) with species specific parameters 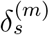 and 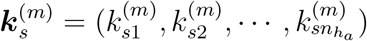 sampled from 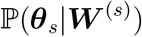.

##### Overlap coefficient Δ_S_

For a given number *n_S_* of sites reflecting the habitat in a region, we sample 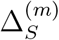 from the posterior distribution 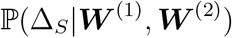 as

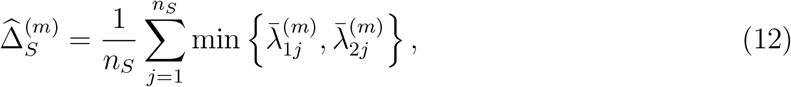

where 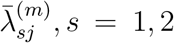 is computed according to equation (3) and normalized such as 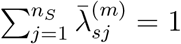 with species specific parameters 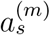 and 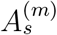 sampled from 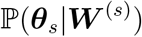.

##### Overlap coefficient Δ_ST_

For *n_T_* time values and *n_J_* number of sites, we sample 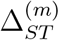 from the posterior distribution 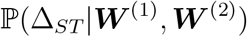 as

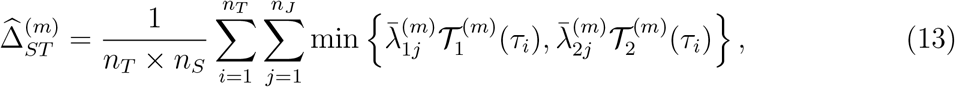

where for species *s* = 1, 2 we calculate 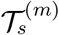 according to equation (6) and 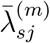 according to equation (3) with species specific parameters 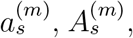 and 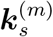 sampled from 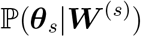, but normalized such that

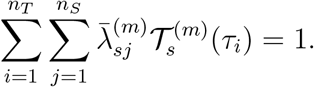

Note that Δ_*ST*_ ≤ Δ_*T*_ and Δ_*ST*_ ≤ Δ_*S*_.

#### 2.1.6. Implementation

All methods were implemented in the `C++` program `Tomcat`, available through a git repository at https://bitbucket.org/WegmannLab/tomcat/ together with a wiki detailing it usage. As input, it requires three files: 1) A description of all camera trap sites in terms of both their start and end date D_*j*1_ and D_*j*2_, respectively, as well as their environmental covariates *X_j_*. 2) A similar file listing the environmental covariates *Y_j_* for each camera trap site. 3) A file containing all observations of a species, consisting of a camera trap identifier and a timestamp. From this input, `Tomcat` will first calculate the full data matrix ***W*** based on the specific choice of *n_o_*.

### 2.2. Simulations

To assess the performance of our algorithm, we generated three sets of simulations:

**Set 1**: To assess the performance of inferring 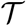, we chose *n_a_* = 24 intervals of *h_a_* = 1h each with *k*_1:4_ = *k*_13:16_ = (1 + *α*, 1 + 3*α*, 1 + 3*α*, 1 + *α*) and all other *k*_5:12,17:24_ = 1 − *α*. We then used *α* = 0, 0.5,1, with *α* = 0 resulting in a uniform activity pattern and *α* = 1 in an activity pattern with two pronounced peaks. We simulated a single site with *a* = 0 and no environmental covariate such that we expect 0.5 observations per day.
**Set 2**: To assess the performance of identifying the correct environmental variables and inferring 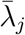, we simulated 50 environmental covariates *X* and *Y*, of which ten each affected 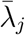 and *p_j_*, respectively. In all cases, we set *A_i_,B_i_* = 0 for all covariates with *γ_λ_i__* = 0 and *γ_pi_* = 0 and set 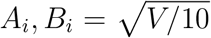 for all covariates with *γ_λ__i_* = 1 and *γ_p_i__* = 1, such that the total variation in 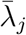 was *V* = 0.1, 0.2, 0.5, 1,2. For these simulations, we set all *k_i_* = 1 and *a* = 0 such that we expect 0.5 observations per day.
**Set 3**: To assess the power of inferring overlap coefficients, we generated simulations with a single environmental variable for two species *s* = 1,2 with no covariate *Y* and ***k**_s_, A_s_* and *a_s_* chosen to result in the desired overlap coefficients. When simulating under a specific Δ_*T*_, we set *A*_1_ = *A*_2_ = 0, *a*_1_ = *a*_2_ = 0 and chose ***k***_1_ and ***k***_2_ as described under Set 1 but with ***k***_2_ shifted by 6h compared to ***k***_1_. We then used *α* = 0.8, 0.5, 0.2 to obtain Δ_*T*_ = 0.2, 0.5, 0.8. To simulate under a specific Δ_*S*_, we set all ***k***_1_ = ***k***_2_ = 1 and *A*_1_ = 0.5 and then determined the parameters *A*_2_, *a*_1_ and *a*_2_ to match Δ_*S*_ = 0.2, 0.5, 0.8 as described in the supplementary information. To simulate under a specific Δ_*ST*_, we set *A*_1_ = 0.5 and ***k***_1_ and ***k***_2_ as when simulating under a specific Δ_*T*_, and then determined the parameters *a*_1_, *a*_2_ and *A*_2_ to match Δ_*ST*_ = 0.2, 0.5, 0.8 as described in the supplementary information. For these cases, we used *α* = 0.4, 0.25, 0.1 to give equal weight to the spatial and temporal components, i.e 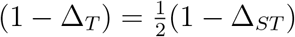.

In all sets, we generated simulations for different total numbers of camera trapping days between 100 and 20,000 with 100 replicates each. For cases with Set 1 and Set 3 with variation in 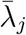, we used *J* = 100 or *J* = 1,000 sites and chose *D* accordingly, excluding combinations for which camera traps were run for less than a full day.

### 2.3. Application to Central African duikers

We applied `Tomcat` to camera trapping data obtained during the dry seasons from 2012 to 2018 from a region in the eastern Central African Republic (CAR), a wilderness exceeding 100,000 km^2^ without permanent settlements, agriculture or commercial logging (Aebischer et al., 2017, 2020). The available data was from 1,059 camera traps set at 532 distinct locations that cover the Aire de Conservation de Chinko (ACC), a protected area of about 20,000 km^2^. For more information about camera deployment and sampling design, see Aebischer et al. (2017). Here, we focus on duikers *Cephalophinae*, which are a diverse mammalian group common in the data set and for which near-perfect manual annotation was available. We set *n_o_* = 15min and used *n_a_* = 24 except for the two species with rather limited data (*Cephalophus leucogaster arrhenii*, and *Cephalophus nigrifrons*).

To infer habitat preferences for these species, we benefited from an existing land cover classification at a 30m resolution that represents the five major habitat types of the Chinko region: moist closed canopy forest (CCF), open savanna woodland (OSW), dry lakéré grassland (DLG), wet marshy grassland (WMG) and surface water (SWA) (Aebischer et al., 2017). Around every camera trap location, we calculated the percentage of each of these habitats in 11 buffers of sizes 30; 65; 125; 180; 400; 565; 1260; 1785; 3,990; 5,640 and 17,840 meters. We complemented this information with the average value within every buffer for each of 15 additional environmental and bioclimatic covariates from the WorldClim database version 2 (Table S.1 Fick and Hijmans, 2017) that we obtained at a resolution of 30 seconds, which translates into a spatial resolution of roughly 1km^2^ per grid cell.

The sparse priors implemented in `Tomcat` allows for the simultaneous use of any number of potentially correlated covariates. To aid in the interpretation of habitat requirements, however, we processed our environmental data as follows: First, we kept only the additional effect of each covariates after regressing out the habitat covariates CCF and OSW at the same buffer. This allows for a direct comparison of the inclusion probabilities of CCF and OSW, since no other covariates may serve as their proxies. Second, we kept only the additional effect of every covariate after regressing out the information contained in the same covariate but at smaller buffers. This ensured that larger buffers may not serve as proxies for smaller ones, and their inclusion in a model implies their importance at that buffer size (see Supporting Information for details).

To predict habitat preferences in the ACC and to calculate Δ_S_ and *Δ_ST_* between species, we determined the same habitat variables at 10,200 regular grid points spaced 2.5 km apart and spanning the entire ACC. To avoid extrapolation, we then restricted our analyses to the 2,639 grid locations that exhibited similar environments to those at which camera traps were placed as measured by the Mahanalobis distance between each grid point and the average across all camera trap locations (see Supporting Information for details).

To characterize detection probabilities, we used the binary classification of the four most common habitat types (CCF, OSW, MWG, DLG) at every location and determined the presence or absence of six additional habitat characteristics: animal path, road, salt lick, mud hole, riverine zone and bonanza.

## 3. Results

### 3.1. Performance against simulations

We first used simulations to assess the performance of `Tomcat` to infer daily activity patterns 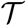 for patterns of different complexity. For each simulation, we quantified the estimation accuracy by calculating 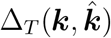 between the true values used in the simulations (**k**) and those inferred 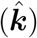. As shown in Figure 2A, 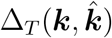 converges towards 1 with increasing data and reached 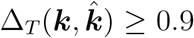 in all cases as of 2,000 camera trapping days, corresponding to about 1,000 observations in these simulations.

**Figure 2:**
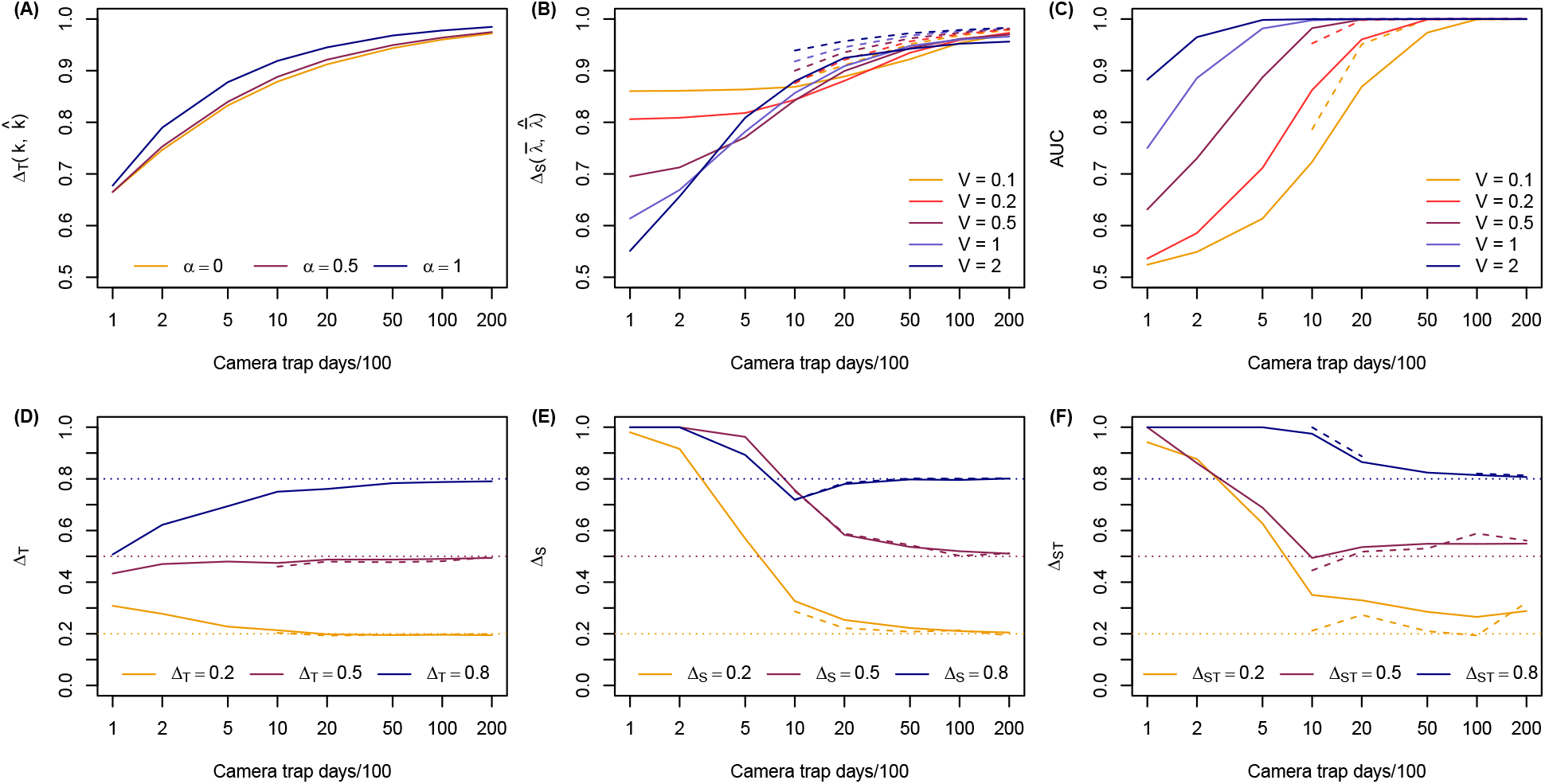
Simulation results as a function of camera trapping days with 0.5 observations per day on average at 100 (solid) or 1,000 sites (dashed). Shown are always averages across 100 replicates. (A) Differences between simulated (***k***) and estimated 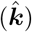 activity patterns as quantified by the overlap 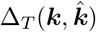 for different values of α. (B) Differences between simulated 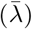 and estimated 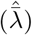 average rates as quantified by the overlap coefficient 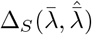 for different values for different values of *V*, the total variance in 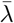. (C) Power to identify environmental covariates as quantified by the Area under the ROC curve (AUC) for the same simulations as in (B). (D-F) Posterior means of inferred overlap coefficients for different true values (dotted lines).

We next used similar simulations to assess the performance of `Tomcat` in inferring the spatial rates 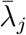, again quantifying the differences between the true 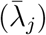 and estimates values (posterior medians 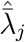) by calculating 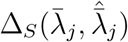. As shown in Figure 2B, the accuracy of the inference increased with more camera trapping days or more sites surveyed. 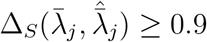 was again reached at a total of 2,000 camera trapping days when 1,000 sites where surveyed (2 days per site), regardless of the effect of environmental variables. If only 100 sites were surveyed, a total of 5,000 camera trapping days (50 days per site) was required to reach this threshold.

Interestingly, the effect of the total variation in 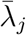 (quantified by *V*) on the estimation accuracy varied as a function of data availability: If data was limited, the inference was less accurate in case *V* was large. If data was more abundant, the inference was more accurate in case *V* was large. This is probably best explained by the power to properly identify contributing environmental covariates, which we quantified by the Area under the ROC Curve (AUC). As shown in Figure 2C and Supplementary Figure S.2, this power increased with more camera trapping days, more sites surveyed, as well as a higher total variance *V*. If data was abundant, high *V* thus resulted in a more accurate identification of the contributing environmental covariates, and therefore in more accurate estimates of 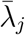. If data was limited, the identification of contributing environmental covariates was more challenging, but errors in this identification had less of an impact if *V* was small.

We further conducted simulations of two species to assess the accuracy in inferring overlap coefficients between species. As shown in Figure 2D-F, all overlap coefficients were accurately inferred as of about 2,000 camera trapping days, with Δ_*T*_ generally requiring the less data than Δ_*S*_ and Δ_*ST*_, in line with the previous findings that ***k*** was inferred more accurately than 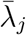. If less than 1,000 camera trapping days (i.e. less than about 500 observations) were used, however, estimates of overlap coefficients were biased towards prior expectations, which are 0.5 and 1.0 for Δ_*T*_ and Δ_*S*_, respectively.

### 3.2. Application to Central African duikers

We used `Tomcat` on existing camera trapping data (Aebischer et al., 2017, 2020) to study the spatio-temporal distribution and overlap of duikers *Cephalophinae* in the Aire de Conservation de Chinko (ACC), a protected area of about 20,000 km^2^ eastern Central African Republic (CAR) spanning the entire savanna-rainforest ecotone (Boulvert, 1985; Olson and Dinerstein, 1998). Duikers are common in the data set and often observed in sympatry, i.e. several species were captured by the same camera trap within a few hours. We detected a total of eight species in the data set (Table 1): Eastern Bay Duiker *Cephalophus dorsalis castaneus*, Uele White Bellied Duiker *Cephalophus leucogaster arrhenii*, Black Fronted Duiker *Cephalophus nigrifrons*, Red Flanked Duiker *Cephalophus rufilatus*, Western Yellow Backed Duiker *Cephalophus silvicultor castaneus*, Weyns Duiker *Cephalophus weynsi*, Eastern Blue Duiker *Philantomba monticola aequatorialis* and Bush Duiker *Sylvicapra grimmia*.

**Table 1:** Available data on the eight detected species of duikers.

| **Species** | **Common Name** | **Pictures** | **Camera Traps** |
| --- | --- | --- | --- |
| *Cephalophus dorsalis castaneus* | Eastern Bay Duiker | 1,631 | 66 |
| *Cephalophus leucogaster arrhenii* | Uele White Bellied Duiker | 432 | 11 |
| *Cephalophus nigrifrons* | Black Fronted Duiker | 102 | 7 |
| *Cephalophus rufilatus* | Red Flanked Duiker | 5,762 | 168 |
| *Cephalophus silvicultor castaneus* | Western Yellow Backed Duiker | 10,321 | 222 |
| *Cephalophus weynsi* | Weyns Duiker | 10,037 | 146 |
| *Philantomba monticola aequatorialis* | Eastern Blue Duiker | 50,979 | 212 |
| *Sylvicapra grimmia* | Bush duiker | 5,626 | 124 |

The eight duiker species varied greatly in their habitat preferences (Figures 3, S.5) as inferred by `Tomcat`. As shown in Figure 3, *C. dorsalis* and *C. weynsi* both have a strong preference for CCF over OSW habitat at the smallest buffers, in contrast to *S. grimmia* that shows a strong preference for OSW. At higher buffers the signal is less clear, probably owing to the heterogeneous nature of the study area, in which both CCF and OSW correlated negatively with WMG and DLG, habitats not well suited for any of these species. Interestingly, *P. monticola* and *C. silvicultor* seem to be true ecotone species preferring a mixture of the canonical habitats CCF and OSW (Figure S.5).

**Figure 3:**
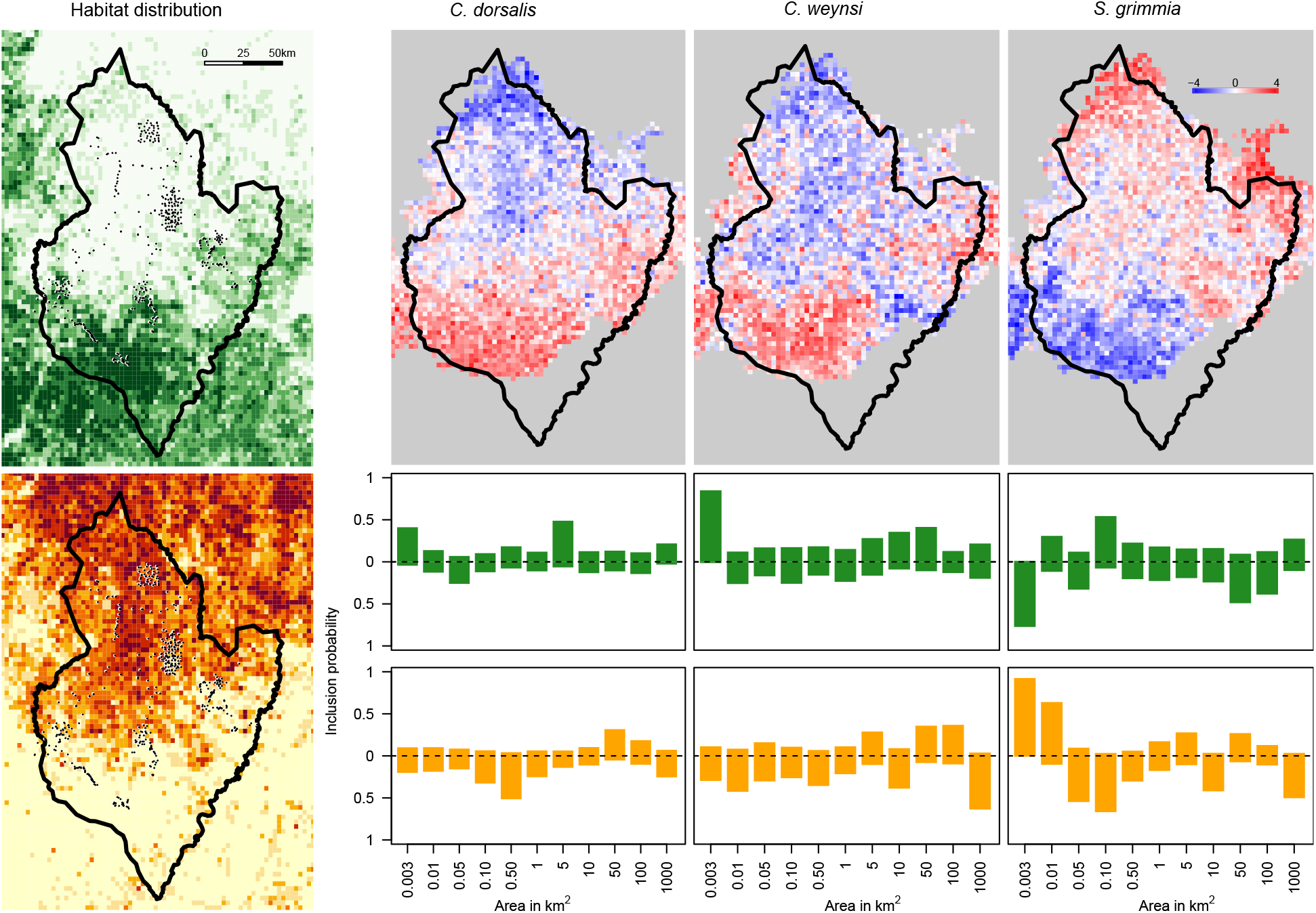
Habitat preference of the three duiker species *C. dorsalis, C. weynsi* and *S. grimmia*. Left: Distribution of closed canopy forest (CCF, top, green) and open savanna woodland (OSW, bottom, yellow) across the study region with the ACC borders and camera trap locations (black dots). Top right: Relative densities *d_sj_* of the three duikers predicted at 2,639 grid points. For each species the colors indicates 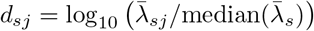, where median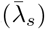 is the median value over all the grid points *j*. Red shades indicate *d_sj_* > 0, blue shades *d_sj_* < 0. Bottom right: Posterior inclusion probabilities for the CCF (green) and OSW (yellow) habitat covariates for each buffer. Values above the dashed line indicate the posterior probability that the habitat correlates positively with the relative species density, values below the dashed line imply a negative correlation.

As shown in Figure 4 and S.4, the species also varied greatly in their daily activity patterns, with some being almost exclusively nocturnal (*C. dorsalis* and *C. silvicultor*), some almost exclusively diurnal (*C. leucogaster, P. monticola, C. nigrifons, C. rufilatus, C. weynsi*) and one crepuscular (*S. grimmia*).

**Figure 4:**
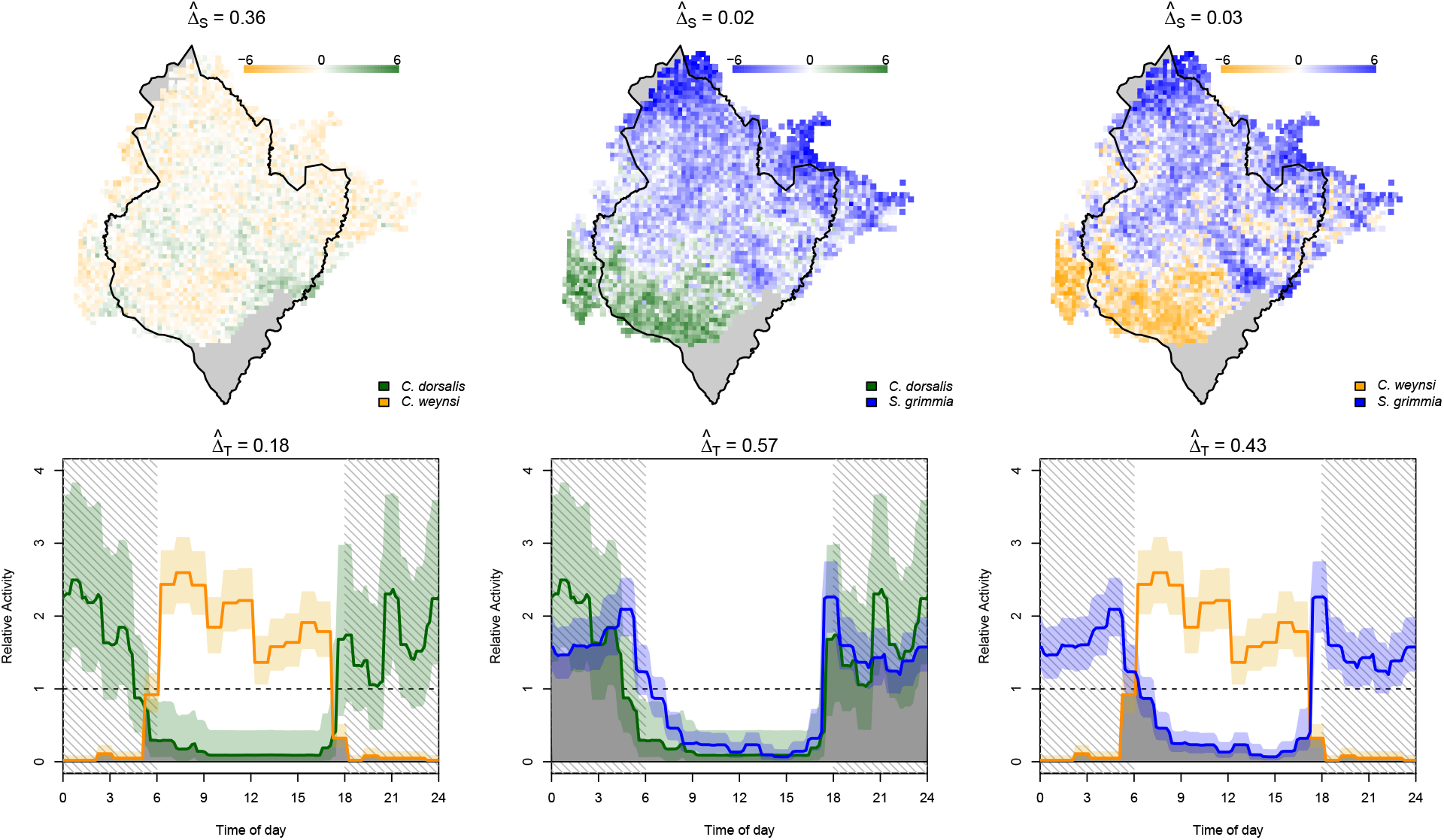
Co-occurrence in space and time between the duiker species *C. dorsalis, C. weynsi*, and *S. grimma*. Top row: interactions in space quantified as 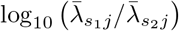 between species 1 and 2. Bottom: posterior mean (solid line) and 90% credible intervals (shades) of temporal activity patterns. The area shaded in gray represents the overlap coefficient Δ_*T*_.

The effect of covariates on detection probabilities followed general expectations: detection probabilities of camera traps placed in CCF or along rivers were estimated as generally lower and those at salt licks and other bonanza as generally higher than average (Supplementary Figure S.3). However, there was considerable variation among species in which covariates impacted detection probabilities, mostly as a result of habitat preferences: for species generally absent at camera traps with a particular covariate (e.g. CCF for savanna species), that covariate was not considered relevant in explaining variation in detection probabilities among observations.

To better understand how these closely related duiker species of similar size and nutritional needs can occur sympatrically, we estimated pairwise overlap coefficients in space and time (Figure 4, Table S.2). Not surprisingly, most species pairs differed substantially either in their habitat preferences or daily activity patterns. Of the two forest dwellers *C. dorsalis* and *C. weynsi* 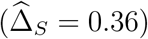 for instance, one is almost exclusively nocturnal and the other almost exclusively diurnal 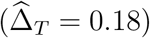, resulting in a small overlap in space and time 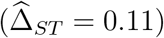. Similarly, the nocturnal *C. dorsalis* and the crepuscular *S. grimmia* that share a lot of temporal overlap 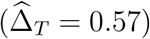 use highly dissimilar habitats 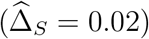, resulting in a very small overlap in time and space 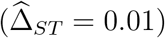.

A visualization using Multidimensional Scaling (MDS) of the pair-wise overlap coefficients of all six species with observations from at least 50 independent camera trap locations is shown in Figure 5. For these species, 84.6% of variation in the temporal overlap can be explained by a single axis separating nocturnal from diurnal species. In contrast, only 44.5% of the variation in the spatial overlap is explained by the first axis distinguishing forest dwellers from savanna species. When using both temporal and spatial information, the two first axis explain 32.3% and 25.5%, respectively, suggesting that a single axis is not sufficient to explain both temporal and spatial differences between species.

**Figure 5:**
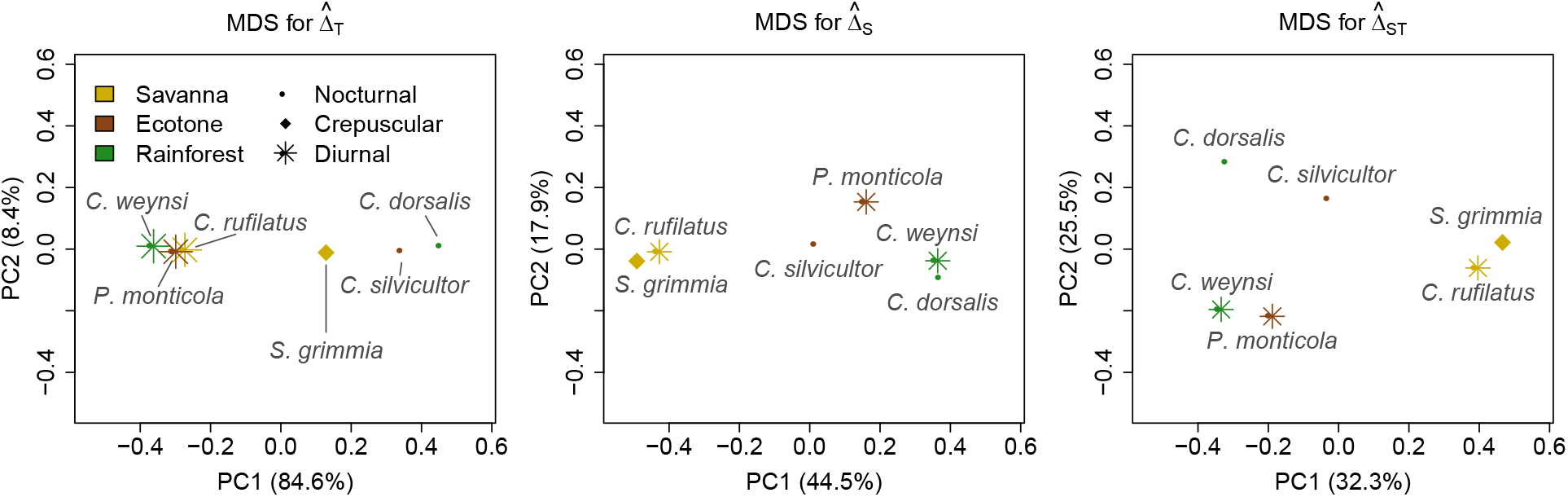
Illustration of the overlap coefficients in time and space between six duiker species visualized in two dimensions using the multidimensional scaling. 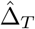: overlap coefficient in time, 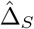: overlap coefficient in space, 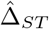: overlap coefficient in time and space.

## 4. Discussion

Devices that continuously record animal observations now make it possible to survey biodiversity of larger or highly vocal animals in relatively short time and with a reasonable budget (Sollmann, 2018; Burton et al., 2015; O’Brien et al., 2010). Thanks to increased battery life, larger media to store data and other technical advances, such data sets can now be produced with comparatively little manual labor, even under the demanding conditions of large and remote areas. In addition, the annotation of these data sets on the species level is now aided by machine learning algorithms that automatize the detection of common species and recordings without observations (e.g. Norouzzadeh et al., 2018). As a result, continuous recording devices have become an indispensable tool for wildlife monitoring.

Of particular interest is the inference of the spatial distribution of species. Traditionally, such distributions are inferred with occupancy models that account for the variability in detection rates between surveyed locations, addressing a key feature of data gathered through ecological surveys (MacKenzie et al., 2002; Sollmann, 2018; Burton et al., 2015). However, the major draw-back of occupancy is that the species distribution is represented as a simple presence-absence matrix not reflective of differences in abundance at occupied sites.

To address this concern, we propose here to decompose the rate of records Λ_*j*_(*t*) of a species at site *j* and time *t* (e.g. the photographic rate) into three components: a spatial component 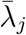 reflecting the average rate of observations at location *j*, a temporal component 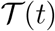 reflecting the daily activity pattern, and the location-specific detection rate *p_j_*. To ensure identifiability, and similar to occupancy models, we further assume that the spatial component 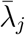 and the detection rates *p_j_* are functions of location-specific covariates (e.g. the environment).

The interpretation of the temporal component 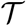 is straight forward and matches that of other methods used to infer daily activity patterns from such data (Ridout and Linkie, 2009). The interpretation of the spatial component 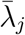 and the detection probabilities *p_j_* warrant some discussion as the model has no intrinsic way to distinguish between them: it is their product 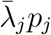 that describes the rate of observations, which itself is affected by the local abundance and activity of the studied species.

In occupancy models, the variation in the rates of observations at occupied sites is attributed to differences in detection rates. Hence, sites at which a species is more abundant or more active will result in higher detection rates. To infer abundances, it is therefor usually assumed that activity does not vary between sites. Royle, J. Andrew and Nichols (2003), for instance, introduced an important extension of occupancy models that infers local abundances from variation in detection rates. In their model, the detection probability is a function of the detection probability of a single individual r and the local abundance *N_j_*. To infer *N_j_* or its distribution, r is then assumed not to vary among sites, implying constant activity.

Most established survey methods that rely on direct or indirect observations to quantify abundances make similar assumptions. Inferring local abundances from responses at call-up stations, for instance, requires knowledge on local response rates (Webster et al., 2010). Similarly, inferring local abundances from observations on transects (Buckland et al., 2001) requires knowledge on local daily travels distances, rates of nest building, or similar quantities depending in the nature of the observation. In practice, estimates of such quantities are at best assessed for a handful of locations (e.g. Funston et al., 2010), but usually borrowed from other studies (e.g. Aebischer et al., 2020; Mathewson et al., 2008).

In `Tomcat`, the variation in the rates of observations may be attributed to either 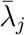 or *p_j_*. By estimating *p_j_* jointly with 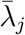, `Tomcat` explicitly accounts for the variation in detection between sites, also that caused by variation in activity. If this variation is captured well, 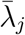 may serve as a good surrogate for the variation in abundance between sites. But since 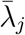 and *p_j_* are confounded, their identification depends on the covariates *X* and *Y* used: if a covariate in *Y* explains part of the variation in the rate of observations, it is included in the model and contributes to *p_j_*. If that covariate was included in *X* instead, it would contribute to 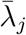. The choice of covariates therefore determines which effects we wish to interpret as underlying 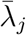 (abundance) and which as underlying *p_j_* (detection or activity).

In the application to duikers, we chose to use the covariates describing the environment (e.g. the habitat or humidity) as relevant for 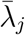 and those describing local features (e.g. the presence of a salt lick or road) as relevant for *p_j_*. The motivation for this choice was our interest to learn about the environmental covariates explaining variation in abundance in terms of 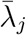, while explaining high photographic rates due to increased activity through pj. However, and depending on the question, other choices may be equally interesting. If the surveyed area is small in relation to mobility, for instance, individuals are likely observed at many sites and 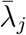 will reflect habitat use of these individuals.

### 4.1. Underlying assumptions

As mentioned above, the `Tomcat` model does not make any assumption about site closure. However, it does assume that records are independent between surveyed sites. That does not imply that a single individual may not be recorded regularly at multiple sites, but it implies that the times at which an individual is recorded at different sites is independent given the general activity pattern 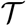 of that species. For instance, two camera traps in close proximity along a deer crossing will not be independent as a record at one is mostly followed by a record at the other. Such records do not provide independent information about the importance of environmental covariates.

In its current implementation, `Tomcat` further assumes that while 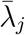 and *p_j_* vary spatially (as functions of environmental covariates), all temporal variation is captured by the daily activity patterns 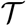. However, detection probabilities may also vary throughout the day, as it might be harder (or easier), for instance, to identify a certain species on black-and-white camera trap pictures taken at night or on infrared pictures taken during cooler times. In its current implementation, the model would explain such variation through 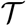, which might lead to biases. An even stronger temporal assumption is that neither 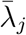, *p_j_* nor 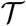 changes seasonally. However, the model could be readily extended to multiple seasons. In the occupancy framework, this is commonly done by modeling extinctions and colonizations explicitly (MacKenzie et al., 2003). In the framework proposed here, an analogue would be to model trends in 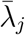 over time. A simple extension would be to model 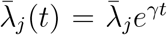 to capture general population trends. To capture seasonal variation over the year, one could modulate 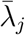 with an additional piecewise-constant function similar to 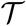 spanning an entire year. Similar extensions can easily be envisioned for the detection probabilities *p_j_* or activity patterns 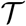.

### 4.2. Species interactions

Records from camera traps or other continuous recording devices may also be used to study the interaction of species in space and time. To infer temporal interactions, two classes of methods exist (Niedballa et al., 2019): In a first class, temporal avoidance is quantified as the degree to which a first species influences subsequent visits of a second species (Harmsen et al., 2009; Karanth et al., 2017). These methods generally compare time intervals between observations of the first and the second species to determine statistical dependence. In a second class, daily activity patterns are inferred individually for each species and then compared using overlap coefficients (Ridout and Linkie, 2009).

Tomcat implements this second class by estimating overlap coefficients between activity patterns 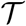 individually inferred for each species. We further extend the concept of overlap to space by comparing the spatial distributions of two species as captured by their respective 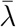 predictions in a specific region. Benefiting from a joint estimation of 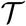 and 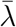, `Tomcat` also estimates overlap coefficients in the spatio-temporal distribution. As we show with simulations, these overlap coefficients can be inferred rather accurately if several hundred observations per species have been recorded.

If fewer observations are present, they are biased towards their prior expectations. This is particularly apparent for Δ_*T*_ that is biased away from 1.0 and may hence result in wrongly inferred differences between species. Niedballa et al. (2019) previously reported this issue and proposed a bootstrap approach to test if an estimate of Δ_*T*_ is significantly different from 1.0.

Just as temporal overlaps, the spatial overlap estimated by `Tomcat` is estimated based on spatial distributions inferred for each species individually. A more powerful approach has been proposed for occupancy models in which spatial interactions are inferred through the probabilities if joint occupancy in multi-species models (MacKenzie et al., 2004; Rota et al., 2016). While not currently implemented, a similar extension can be envision for the model proposed here by adding an interaction term to the 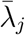.

### 4.3. Co-existence of Duikers in a Savanna-rainforest ecotone

We here used `Tomcat` to infer the spatio-temporal distribution of eight species of duikers sympatrically occurring in the Aire de Conservation de Chinko (ACC) in the Central African Republic. These species varied greatly both in their daily activity patterns and habitat preferences, with some species being almost exclusively nocturnal, others diurnal or crepuscular. Similar, some species showed a strong preference for close canopy forest (CCF), some for open savanna woodland (OSW), and two appeared to be true ecotone species with a preference for mixed habitat. An interesting observation was that the two rarely studied species *C. weynsi* and *C. leucogaster* not only occur in large blocks of CCF as suggested in the literature (Kingdon et al., 2013), but also in narrow gallery forests within the forest-savanna ecotone several kilometers away from the next extensive forest block (Figures 3 and S.5).

When comparing these species using spatio-temporal overlap coefficients, we found that frequently observed and therefore evidently abundant species within this community tend to differ in their habitat preference and/or daily activity. In contrast, infrequently observed and therefore putatively rare taxa seem to have large overlap with co-occurring species. *C. leucogaster*, for instance, which is rather rare and was only observed at 11 distinct locations (Table 1), has similar habitat preferences and is active at the same time as *C. weynsi*, which is among the most common forest duikers within the ACC. In contrast, *C. dorsalis*, which is strictly nocturnal, seems to co-exist with the *C. weynsi* at higher densities (Figure 4, Table S.2).

### 4.4. Conclusion

We here introduced `Tomcat`, a model that infers habitat preferences and daily activities from imperfect spatio-temporal observations. Similar to occupancy models, it allows to learn about the ecological requirements of animals, including rare, elusive and unmarked species. But unlike occupancy models, it does not estimate the presence or absence of a species, but rather a measure informative about species densities under the mild assumptions that the jointly modeled detection probabilities capture variation in activity between sites. In addition, `Tomcat` explicitly models daily activity patterns. While estimating these quantities may require larger data sets than the inference under occupancy models, we believe they constitute a major step forward in understanding and monitoring species distributions.

## Acknowledgments

We thank two anonymous reviewers for their exceptionally detailed and constructive feed-back. This work was supported by a Swiss National Science Foundation grant (31003A_173062) to DW.

## Author contributions

TA and DW conceived the idea; SA, BW, MC and DW developed and implemented the method; TA collected the data; SA, LS, TA and MC analyzed the data; SA, LS and DW led the writing of the manuscript. All authors contributed critically to the drafts and gave final approval for publication.

## References

Aebischer, T., Ibrahim, T., Hickisch, R., Furrer, R.D., Leuenberger, C., Wegmann, D., 2020. Apex predators decline after an influx of pastoralists in former Central African Republic hunting zones. Biological Conservation 241. doi:10.1016/j.biocon.2019.108326.

Aebischer, T., Siguindo, G., Rochat, E., 2017. First quantitative survey delineates the distribution of chimpanzees in the eastern central african republic. Biological Conservation 213, 84–94.

Boulvert, Y., 1985. Carte phytogeographique de la republique centrafricaine. ORSTOM (Office de la recherche scientifique et technique Outre-Mer).

Bu, H., Wang, F., Mcshea, W.J., Lu, Z., Wang, D., Li, S., 2016. Spatial Co-Occurrence and Activity Patterns of Mesocarnivores in the Temperate Forests of Southwest China. PLOS ONE, 1–15doi:10.1371/journal.pone.0164271.

Buckland, S.T., Anderson, D.R., Burnham, K.P., Laake, J.L., Borchers, D.L., Thomas, L., et al., 2001. Introduction to distance sampling: estimating abundance of biological populations.

Burton, A.C., Neilson, E., Moreira, D., Ladle, A., Steenweg, R., Fisher, J.T., Bayne, E., Boutin, S., 2015. Review: Wildlife camera trapping: a review and recommendations for linking surveys to ecological processes. Journal of Applied Ecology 52, 675–685. doi:10.1111/1365-2664.12432.

Caravaggi, A., Banks, P.B., Burton, A.C., Finlay, C.M.V., Haswell, P.M., Hayward, M.W., Rowcliffe, M.J., Wood, M.D., 2017. A review of camera trapping for conservation behaviour research. Remote Sensing in Ecology and Conservation 3, 109–122. doi:10.1002/rse2.48.

Efford, M.G., Dawson, D.K., 2012. Occupancy in continuous habitat. Ecosphere 3, art32. doi:10.1890/ES11-00308.1.

Fick, S.E., Hijmans, R.J., 2017. Worldclim 2: new 1-km spatial resolution climate surfaces for global land areas. International Journal of Climatology 37, 4302–4315. doi:10.1002/joc.5086.

Frey, S., Fisher, J.T., Burton, A.C., Volpe, J.P., 2017. Investigating animal activity patterns and temporal niche partitioning using camera-trap data: challenges and opportunities. Remote Sensing in Ecology and Conservation 3, 123–132. doi:10.1002/rse2.60.

Funston, P.J., Frank, L., Stephens, T., Davidson, Z., Loveridge, A., Macdonald, D., Durant, S., Packer, C., Mosser, A., Ferreira, S.M., 2010. Substrate and species constraints on the use of track incidences to estimate african large carnivore abundance. Journal of Zoology 281, 56–65.

Gelman, A., Rubin, D.B., 1992. Inference from iterative simulation using multiple sequences. Statistical Science 7, 457–472. URL: http://www.jstor.org/stable/2246093.

Harmsen, B.J., Foster, R.J., Silver, S.C., Ostro, L.E.T., Doncaster, C.P., 2009. Spatial and Temporal Interactions of Sympatric Jaguars (Panthera onca) and Pumas (Puma concolor) in a Neotropical Forest. Journal of Mammalogy 90, 612–620. doi:10.1644/08-MAMM-A-140R.1.

Karanth, K.U., Srivathsa, A., Vasudev, D., Puri, M., Parameshwaran, R., Kumar, N.S., 2017. Spatio-temporal interactions facilitate large carnivore sympatry across a resource gradient. Proceedings of the Royal Society B: Biological Sciences 284, 20161860. doi:10.1098/rspb.2016.1860.

Kingdon, J., Happold, D., Butynski, T., Hoffmann, M., Happold, M., Kalina, J., 2013. Mammals of Africa. v. 1-6, Bloomsbury Publishing.

Kriticos, D.J., Jarosik, V., Ota, N., 2014. Extending the suite of bioclim variables: a proposed registry system and case study using principal components analysis. Methods in Ecology and Evolution 5, 956–960.

MacKenzie, D.I., Bailey, L.L., Nichols, J.D., 2004. Investigating species co-occurrence patterns when species are detected imperfectly. Journal of Animal Ecology 73, 461–555. doi:10.1111/j.0021-8790.2004.00828.x.

MacKenzie, D.I., Nichols, J.D., 2004. Occupancy as a surrogate for abundance estimation. Animal biodiversity and conservation 27, 461–467.

MacKenzie, D.I., Nichols, J.D., Hines, J.E., Knutson, M.G., Franklin, A.B., 2003. Estimating site occupancy, colonization, and local extinction when a species is detected imperfectly. Ecology 84, 2200–2207. doi:10.1890/02-3090.

MacKenzie, D.I., Nichols, J.D., Lachman, G.B., Droege, S., Andrew Royle, J., Langtimm, C.A., 2002. Estimating site occupancy rates when detection probabilities are less than one. Ecology 83, 2248–2255. doi:10.1890/0012-9658(2002)083[2248:ESORWD]2.0.CO;2.

MacKenzie, D.I., Royle, J.A., 2005. Designing occupancy studies: general advice and allocating survey effort. Journal of Applied Ecology 42, 1105–1114. doi:10.1111/j.1365-2664.2005.01098.x.

Mathewson, P.D., Spehar, S.N., Meijaard, E., Nardiyono, Purnomo, Sasmirul, A., Sudiyanto, Oman, Sulhnudin, Jasary, Jumali, Marshall, A.J., 2008. Evaluating orangutan census techniques using nest decay rates: Implications for population estimates. Ecological Applications 18, 208–221. doi:10.1890/07-0385.1.

Niedballa, J., Wilting, A., Sollmann, R., Hofer, H., Courtiol, A., 2019. Assessing analytical methods for detecting spatiotemporal interactions between species from camera trapping data. Remote Sensing in Ecology and Conservation 5, 272–285. doi:10.1002/rse2.107.

Norouzzadeh, M.S., Nguyen, A., Kosmala, M., Swanson, A., Palmer, M.S., Packer, C., Clune, J., 2018. Automatically identifying, counting, and describing wild animals in camera-trap images with deep learning. Proceedings of the National Academy of Sciences 115, E5716–E5725. doi:10.1073/pnas.1719367115.

O’Brien, T.G., Baillie, J.E.M., Krueger, L., Cuke, M., 2010. The wildlife picture index: monitoring top trophic levels. Animal Conservation 13, 335–343. doi:10.1111/j.1469-1795.2010.00357.x.

Oliveira-santos, L.G.R., Zucco, C.A., Agostinelli, C., 2013. Using conditional circular kernel density functions to test hypotheses on animal circadian activity. Animal Behaviour 85, 269–280. doi:10.1016/j.anbehav.2012.09.033.

Olson, D.M., Dinerstein, E., 1998. The global 200: A representation approach to conserving the earth’s most biologically valuable ecoregions. Conserv. Biol. 12, 502–515.

Ridout, M.S., Linkie, M., 2009. Estimating overlap of daily activity patterns from camera trap data. Journal of Agricultural, Biological, and Environmental Statistics 14, 322–337. doi: 10.1198/jabes.2009.08038.

Rota, C.T., Ferreira, M.A.R., Kays, R.W., Forrester, T.D., Kalies, E.L., McShea, W.J., Parsons, A.W., Millspaugh, J.J., 2016. A multispecies occupancy model for two or more interacting species. Methods in Ecology and Evolution 7, 1164–1173. doi:10.1111/2041-210X.12587.

Rowcliffe, J.M., Kays, R., Kranstauber, B., Carbone, C., Jansen, P.A., 2014. Quantifying levels of animal activity using camera trap data. Methods in Ecology and Evolution 5, 1170–1179.

Royle, J. Andrew and Nichols, J.D., 2003. Estimating Abundance From Repeated Presence – Absence. Ecology 84, 777–790.

Sollmann, R., 2018. A gentle introduction to camera-trap data analysis. African Journal of Ecology 56, 740–749. doi:10.1111/aje.12557.

Steenweg, R., Hebblewhite, M., Whittington, J., Lukacs, P., McKelvey, K., 2018. Sampling scales define occupancy and underlying occupancy–abundance relationships in animals. Ecology 99, 172–183. doi:10.1002/ecy.2054.

Title, P.O., Bemmels, J.B., 2018. Envirem: an expanded set of bioclimatic and topographic variables increases flexibility and improves performance of ecological niche modeling. Ecography 41, 291–307.

Tobler, M.W., Hartley, A., Carrillo-Percastegui, S.E., Powell, G.V.N., 2015. Spatiotemporal hierarchical modelling of species richness and occupancy using camera trap data. Journal of Applied Ecology 43, 413–421.

Webster, H., McNutt, J.W., McComb, K., 2010. Eavesdropping and risk assessment between lions, spotted hyenas and african wild dogs. Ethology 116, 233–239. doi:10.1111/j.1439-0310.2009.01729.x.

