## Supplementary Information for "A sparse observation model to quantify species interactions in time and space"

—

### Supporting Information

#### S1. Details on MCMC

*Proposal Kernels*

To generate samples from the posterior distribution $\mathbb{P}(\boldsymbol{\theta}|\boldsymbol{W})$, we use an MCMC algorithm with Metropolis-Hastings updates with the following proposal kernels:

- For the update $a \to a'$, we use a symmetric proposal kernel $a' \sim \mathcal{N}(a, \sigma^2_{a\to a'})$.
- For the update $A_i \to A'_i$ we used a symmetric proposal kernel $A'_i \sim \mathcal{N}(A_i, \sigma^2_{A\to A',\gamma_{\lambda_i}})$ with different variances $\sigma^2_{A\to A',0}$ and $\sigma^2_{A\to A',1}$ depending on $\gamma_{\lambda_i}$. The updates $B_i \to B'_i$ are analogous with parameters $\sigma^2_{B\to B',0}$ and $\sigma^2_{B\to B',1}$.
- For the update $\gamma_{\lambda_i} \to \gamma'_{\lambda_i}$, we always propose $\gamma'_{\lambda_i} = 1 - \gamma_{\lambda_i}$, and analogously for the updates $\gamma_{p_i} \to \gamma'_{p_i}$.
- For the update $\sigma_{\lambda 0} \to \sigma'_{\lambda 0}$, we use a symmetric proposal kernels $\sigma'_{\lambda 0} \sim \mathcal{N}(\sigma_{\lambda 0}, \sigma^2_{\sigma_{\lambda 0}\to\sigma'_{\lambda 0}})$, mirrored at 0, and analogously for the updates $\sigma_{\lambda 1} \to \sigma'_{\lambda 1}$, $\sigma_{p0} \to \sigma'_{p0}$ and $\sigma_{p1} \to \sigma'_{p1}$ with parameters $\sigma^2_{\sigma_{\lambda 1}\to\sigma'_{\lambda 1}}$, $\sigma^2_{\sigma_{p0}\to\sigma'_{p0}}$ and $\sigma^2_{\sigma_{p1}\to\sigma'_{p1}}$, respectively.
- For the update $\boldsymbol{k} \to \boldsymbol{k}'$, we begin by picking a random activity interval $i$ and proposing a move $k_i \to k'_i$ according to a symmetric transition kernel $k'_i \sim \mathcal{N}(k_i, \sigma^2_{k\to k'})$ mirrored at 0. We then scale all other entries of $\boldsymbol{k}'$ to satisfy the constraint on the sum. Specifically, we set $k'_j = \alpha k_j$ for all $j \neq i$, where

$$\alpha = \frac{n - k'_i}{\sum_{j\neq i} k_j}.$$

- For the update $\delta \to \delta'$, we choose $\delta' = \delta + 1$ or $\delta' = \delta - 1$ with equal probability, but wrap around at modulus $h_a - h_o$.

- For the update $\pi_\lambda \to \pi'_\lambda$, we use a symmetric proposal kernel $\pi'_\lambda \sim \mathcal{N}(\pi_\lambda, \sigma^2_{\pi_\lambda \to \pi'_\lambda})$ mirrored at 0 and 1. The updates $\pi_p \to \pi'_p$ are analogous with parameter $\sigma^2_{\pi_p \to \pi'_p}$.

We adjust $\sigma^2_{a\to a'}$, $\sigma^2_{A\to A',0}$, $\sigma^2_{A\to A',1}$, $\sigma^2_{B\to B',0}$, $\sigma^2_{B\to B',1}$, $\sigma^2_{\sigma_{\lambda 0}\to\sigma'_{\lambda 0}}$, $\sigma^2_{\sigma_{\lambda 1}\to\sigma'_{\lambda 1}}$, $\sigma^2_{\sigma_{p0}\to\sigma'_{p0}}$, $\sigma^2_{\sigma_{p1}\to\sigma'_{p1}}$, $\sigma^2_{k\to k'}$, $\sigma^2_{\pi_\lambda\to\pi'_\lambda}$ and $\sigma^2_{\pi_p\to\pi'_p}$ during a round of successive burnins. Here, we used ten burnins of 1000 iterations each.

*Initialization*

We initialize the MCMC algorithm as follows:

- We set all $k_i = 1, i = 1, \dots n_{h_a}$.
- We set $\pi_\lambda$ and $\pi_p$ to user provided values.
- We obtain an initial estimate of $a$ from the fraction $f$ of intervals $\boldsymbol{W}$ that have at least one observation. From (5) and assuming that $A = 0$ and all $p_j = \frac{1}{2}$ (see above), we get

$$a = -\frac{2}{h_o}\log(1-f). \tag{S.14}$$

- Starting with a current estimate $A = 0, B = 0$, we iteratively add estimates

$$\hat{A}_i = \arg\max_{A_i} \ \mathcal{L}(a, A_{-i}, A_i, B, \delta, \boldsymbol{k}),$$

  for the covariates maximizing $\mathcal{L}(a, A_{-i}, \hat{A}_i, B, \delta, \boldsymbol{k})$, until a fraction $\pi_\lambda$ of the coefficients are non-zero. Here, $A_{-i}$ denotes the current estimate of $A$ without entry $i$ and $\mathcal{L}(\cdot)$ is given by (7). The same procedure is then used to initialize $B$ until a fraction $\pi_p$ of the coefficients are non-zero.

#### S2. Simulating data with specific overlap coefficients

*Simulating data with specific $\Delta_S$*

We wish to simulate scenarios for which $\mathbb{E}[\Delta_S] = \delta_S$ where the expectation is across covariate $X$. From (12) we get

$$\mathbb{E}[\Delta_S] = \mathbb{E}\left[\min\left\{\bar{\lambda}_1(X), \bar{\lambda}_2(X)\right\}\right] = \int \min\left(\bar{\lambda}_1(X), \bar{\lambda}_2(X)\right)\mathbb{P}(X)\mathrm{d}X \tag{S.15}$$

with constraints

$$\int \bar{\lambda}_i(X)\mathbb{P}(X)\mathrm{d}X = 1; s = 1, 2, \tag{S.16}$$

and $\bar{\lambda}_i$ given by equation (3).

For a single covariate $X \sim \mathcal{N}(0,1)$ and parameters $a_i$ and $A_i$ for species $i$ we thus have

$$\int \exp(a_i + A_i X)\varphi(X)\mathrm{d}X = 1,$$

where $\varphi(\cdot)$ denotes the density of the standard normal distribution. From this it follows that

$$\exp(a_i + \frac{1}{2}A_i^2) = 1 \Leftrightarrow a_i = -\frac{1}{2}A_i^2. \tag{S.17}$$

For a specific draw of the covariate $X = x$ We further have

$$\begin{aligned}\bar{\lambda}_1(x) < \bar{\lambda}_2(x) &\Leftrightarrow \exp(-\tfrac{1}{2}A_1^2 + A_1 x) < \exp(-\tfrac{1}{2}A_2^2 + A_2 x) \\ &\Leftrightarrow x < \tfrac{A_1+A_2}{2} = u_x.\end{aligned}$$

From (S.15) we have

$$\begin{aligned}\mathbb{E}[\Delta_S] &= \int \min\left\{\bar{\lambda}_1(X), \bar{\lambda}_2(X)\right\}\mathbb{P}(X)\mathrm{d}X \\ &= \int_{-\infty}^{u_x} \exp(-\frac{1}{2}A_1^2 + A_1)\varphi(X)\mathrm{d}X + \int_{u_x}^{+\infty} \exp(-\frac{1}{2}A_2^2 + A_2 X)\varphi(X)\mathrm{d}X \\ &= \Phi\left(\frac{1}{2}(A_2 - A_1)\right) + \left(1 - \Phi\left(\frac{1}{2}(A_1 - A_2)\right)\right) \\ &= 2 - 2\Phi\left(\frac{1}{2}(A_1 - A_2)\right),\end{aligned}$$

where $\Phi)\cdot)$ denotes the cumulative distribution function of the standard normal distribution.

For a specific choice of $A_1$, we can therefore simulate data with $\mathbb{E}[\Delta_S] = \delta_S$ with

$$A_2 = a_1 - 2\Phi^{-1}\left(\frac{2-\delta_S}{2}\right)$$

and $a_1$ and $a_2$ given by (S.17).

*Simulating data with specific $\Delta_{ST}$*

We wish to simulate scenarios for which $\mathbb{E}[\Delta_{ST}] = \delta_{ST}$ where the expectation is across covariate $X$. From (13) we get

$$\mathbb{E}[\Delta_{SR}] = \int\int_0^T \min\left\{\bar{\lambda}_1(X)\mathcal{T}_1(\tau), \bar{\lambda}_2(X)\mathcal{T}_2(\tau)\right\}\mathbb{P}(X)\mathrm{d}\tau\mathrm{d}X \tag{S.18}$$

with constraints

$$\int\int_0^T \bar{\lambda}_i(X)\mathcal{T}_i(\tau)\mathbb{P}(X)\mathrm{d}\tau\mathrm{d}X = \mathbb{E}\left[\mathcal{T}_i(\tau)\bar{\lambda}_{ij}(X_j)\right] = \mathbb{E}\left[\mathcal{T}_i(\tau)\right]\mathbb{E}\left[\bar{\lambda}_{ij}(X_j)\right] = \mathbb{E}\left[\bar{\lambda}_{ij}(X_j)\right] = 1,$$

where $\bar{\lambda}_i$ and $\mathcal{T}_i(\tau)$ for species $i = 1, 2$ are given by (3) and (6), respectively, and $\mathbb{E}\left[\mathcal{T}_i(\tau)\right] = 1$ by definition (see 1). From (S.17) we thus have

$$a_i = -\frac{1}{2}A_i^2.$$

For a specific choice of $A_1$, $\mathcal{T}_1$ and $\mathcal{T}_2$, we can thus simulate data with $\mathbb{E}[\Delta_{ST}] = \delta_{ST}$ by numerically identifying the value of $A_2$ such that

$$\delta_{ST} - \frac{1}{n_T \times n_S}\sum_{i=1}^{n_T}\sum_{j=1}^{n_S}\min\left\{\bar{\lambda}_{1j}^{(f)}\mathcal{T}_1^{(f)}(\tau_i), \bar{\lambda}_{2j}^{(f)}\mathcal{T}_2^{(f)}(\tau_i)\right\} = 0. \tag{S.19}$$

We note that since $\Delta_{ST} \leq \Delta_T$ the choice of $\mathcal{T}_1$ and $\mathcal{T}_2$ will influence the maximally possible $\Delta_{ST}$.

#### S3. Decorrelation environmental covariates

While `Tomcat` readily handles correlated environmental covariates, we chose to decorrelate specific covariates to aid in interpretation. Specifically, we processed our environmental data as follow (two steps):

Step 1: A major interest in our application was to study the impact of the prevalence of closed canopy forest ($f$) and savanna ($s$) habitat on species densities. For each scale (buffer) $b$, we therefore regress each environmental covariate $V_{ib}, i \neq f, s$:

$$V_{ib} = \alpha_{fb}V_{fb} + \alpha_{sb}V_{sb} + \epsilon_{ib}, \tag{S.20}$$

where $V_{fb}$ and $V_{sb}$ represents, respectively, the forest and the Savannah habitat for a buffer $b = 1, \cdots, B$, $i = 1, \cdots, n_{env}$, and $\epsilon_{ib}$ is the error from the linear model described by equation (S.20), which captures the information of the covariate $V_{ib}$ independent of $V_{fb}$ and $V_{sb}$ at buffer $b$. We therefore replace the $V_{ib}$ covariates by $\widetilde{V}_{ib} = \epsilon_{ib}$ in our model, but kept $\widetilde{V}_{fb} = V_{fb}$ and $\widetilde{V}_{sb} = V_{sb}$.

Step 2: To evaluate relevant spatial scale of environmental covariates, we also regressed out the larger buffers from the smaller one as:

$$\widetilde{V}_{ib} = \sum_{j=1}^{b-1} \beta_j \widetilde{V}_{i,j} + e_{ib}, \tag{S.21}$$

In the second step, and for a given environmental covariate $\widetilde{V}_{ib}$ at the scale, we give priority to the smaller scales by keeping only the additional explanation by the studied covariate to what we already know by replacing $\widetilde{V}_{ib}$ by $\widetilde{\widetilde{V}}_{ib} = e_{ib}$.

#### S4. Identifying environmental outliers

Let $X$ be the $J \times E$ matrix of $E$ environmental variables at the $J$ locations surveyed. Let $X^*$ be a $n_S \times E$ be a matrix of the same environmental variables at $n_S$ locations at which we wish to predict $\bar{\lambda}_j$ and wish to use to calculate $\Delta_s$ and $\Delta_{ST}$. To avoid unjustified extrapolation, we chose the $n_S$ locations among all locations on a grid across the ACC as those that matched the following condition: We calculated the Mahanalobis distance $D$ of each surveyed location as

$$D^2 = (X - \mu)^T \Sigma^{-1} (X - \mu),$$

where $\mu$ and $\Sigma$ are the center and variance-covariance matrix of $X$. Similarly, we calculated the Mahalanobis distances $D^*$ of each grid point as

$$D^{*2} = (X^* - \mu)^T \Sigma^{-1} (X^* - \mu),$$

using the same $\mu$ and $\Sigma$ learned form $X$. We then only accepted grid points such that $\max(D^*) \leq 2\max(D)$.

#### S5. Supplementary figures

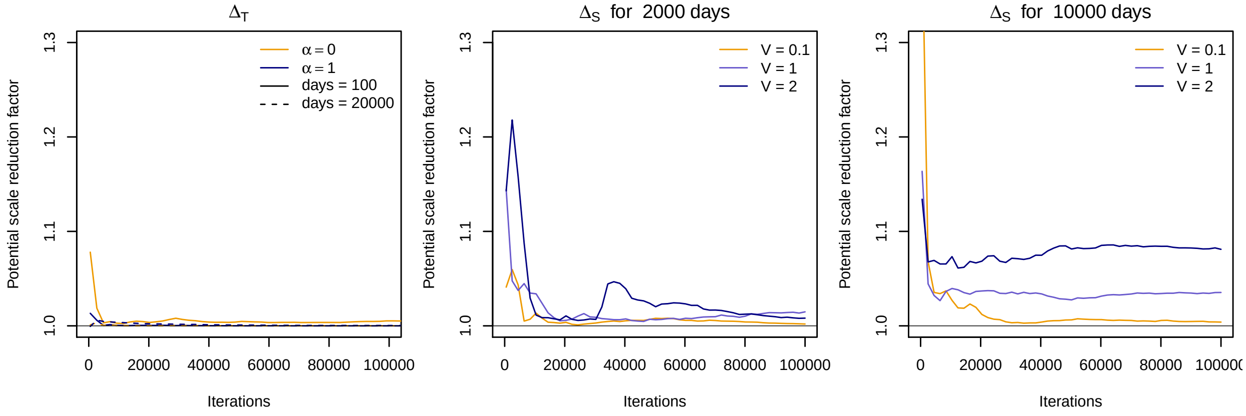


Figure S.1: Convergence of MCMC chains assessed using the Gelman-Rubin diagnostics on five independent runs. Left: Convergence diagnostics for $\Delta_T$ on data simulated at a single site with activity patterns of different complexity ($\alpha = 0$ and $\alpha = 1$) and for limited and extensive data (100 and 20,000 camera trapping days). Middle: Convergence diagnostics for $\Delta_S$ on data simulated at 100 sites surveyed for a total of 2,000 camera trapping days (20 days per site) for different total variances in $\bar{\lambda}_j$ ($V = 0.1$, $V = 1$ and $V = 2$). Right: Same as in Middle but with a total of 10,000 camera trapping days (100 days per site).

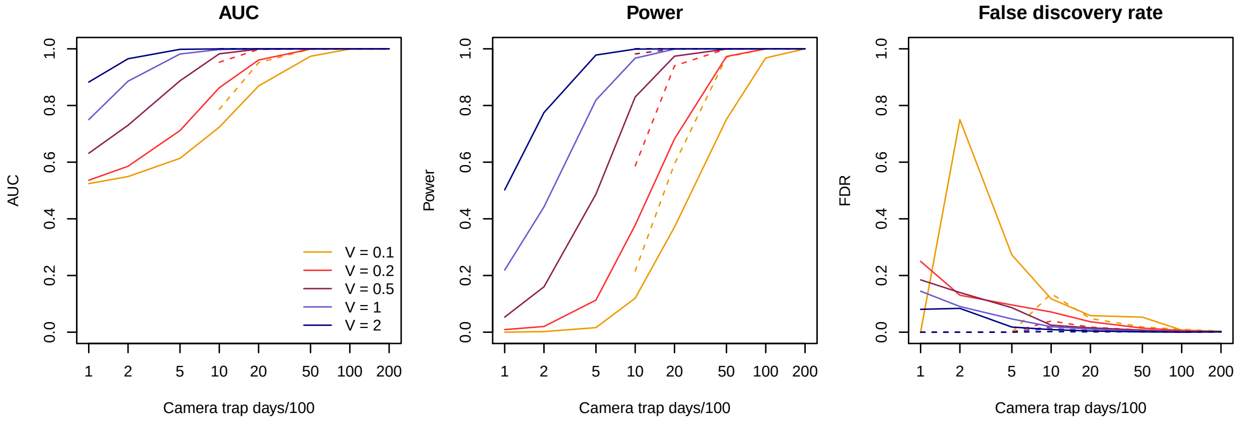


Figure S.2: Power to identify environmental covariates. Shown are the Area under the ROC curve (AUC, left), the power (center) and false discovery rate ar a threshold of 0.9 (right) for the same simulations as in Figure 2B and C. The solid and dashed lines correspond to 100 and 1000 sites, respectively. Small values of the total variance in $\bar{\lambda}_j$, as quantified by $V$, coupled with little data results in high false discovery rates. However, `Tomcat` is still conservative in these cases: for $V = 0.1$, 100 sites and 200 total camera trapping days, only 8 (2 true positives and 6 false positives) of all 5,000 simulated environmental variables were above the inclusion probability threshold of 0.9.

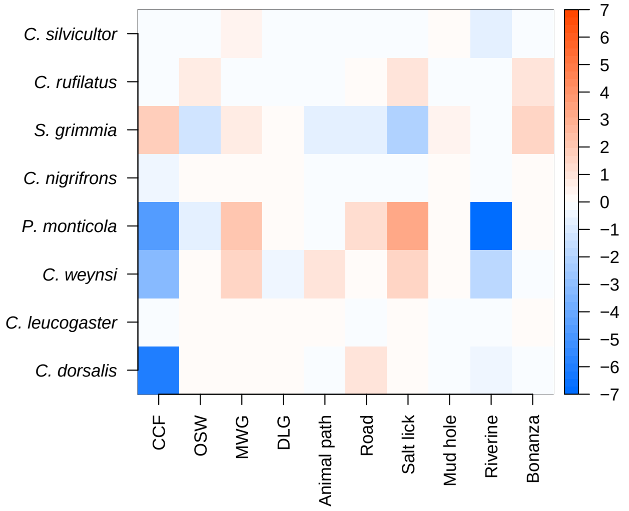


Figure S.3: Effect of covariates on detection probabilities. Shown are the posterior medians of the coefficients $B_i$ for each species (y-axis) and each considered binary covariate $i$ (x-axis). Red and blue shades indicate an increase and decrease in detection probabilities, respectively.

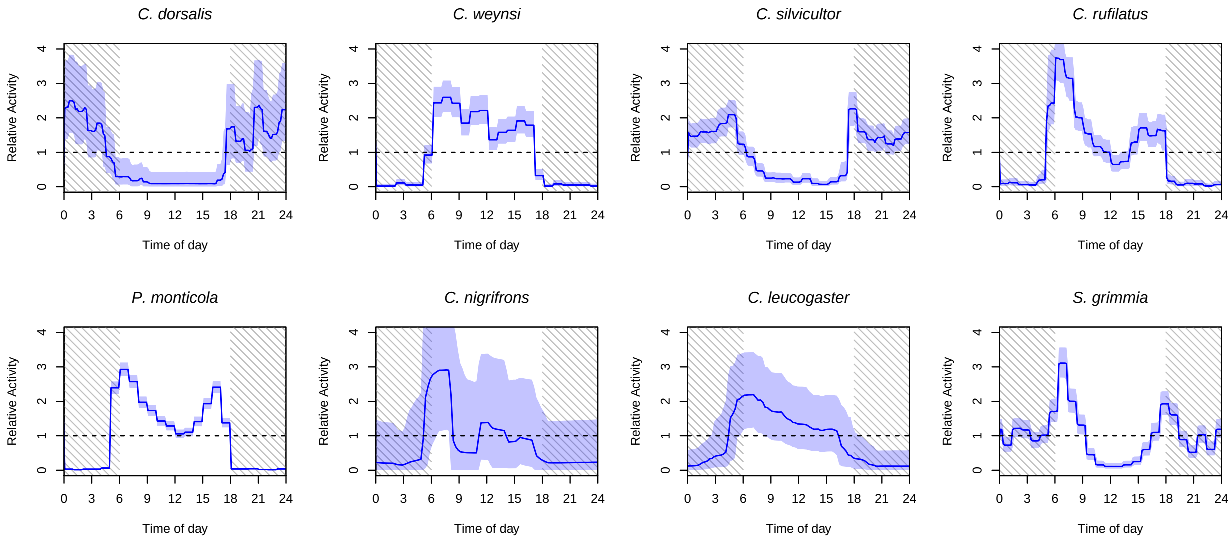


Figure S.4: Estimation of the daily activity of the eight observed duiker species. The solid line represents the posterior mean and the shades, 90% credible intervals of the temporal activity patterns.

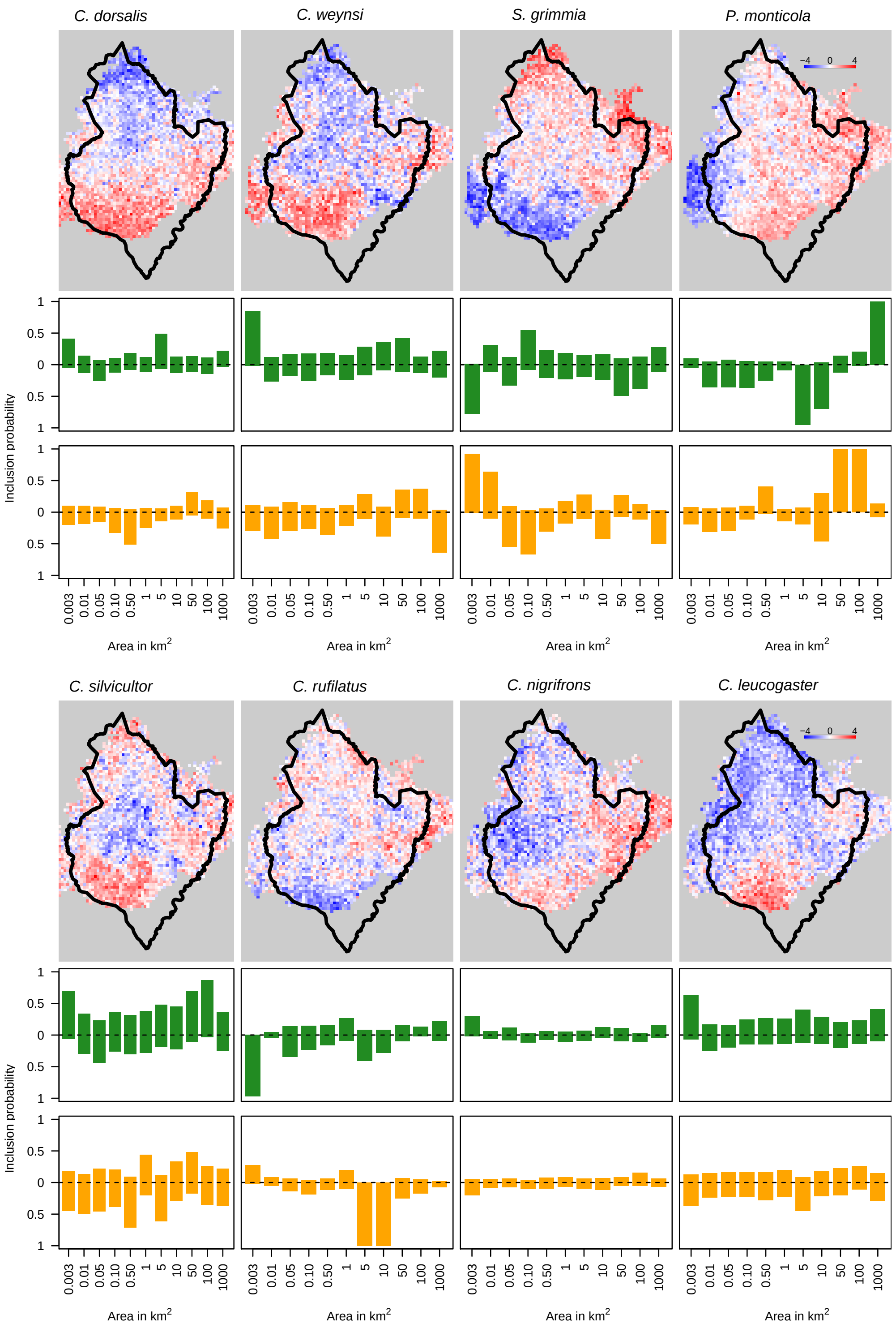


Figure S.5: Relative densities $d_{sj}$ of the eight duiker species predicted at 2,639 grid points. For each species the colors indicates $d_{sj} = \log_{10}\left(\bar{\lambda}_{sj}/\text{median}(\bar{\lambda}_s)\right)$, where median$(\bar{\lambda}_s)$ is the median value over all the grid points $j$. Red shades indicate $d_{sj} > 0$, blue shades $d_{sj} < 0$. Posterior inclusion probabilities for the CCF (green) and OSW (yellow) habitat covariates for each buffer. Values above the dashed line indicate the posterior probability that the habitat correlates positively with the relative species density, values below the dashed line imply a negative correlation.

#### S6. Supplementary tables

Table S.1: Environmental and bioclimatic covariates obtained from the WorldClim database version 2.

| Variable name | Description |
|---|---|
| dem | Digital elevation model |
| slope | Slope of a point |
| aspect | Direction of slope |
| TRI | Terrain Ruggedness Index |
| TWI | Topographic Wetness Index |
| wind | wind speed (m $s^{-1}$) |
| bio2 | Mean Diurnal Range (Mean of monthly (max temp - min temp)) |
| bio5 | Max Temperature of Warmest Month |
| bio6 | Min Temperature of Coldest Month |
| bio12 | Annual Precipitation |
| bio15 | Precipitation Seasonality (Coefficient of Variation) |
| NDVI | Normalized Difference Vegetation Index |
| GC | General Curvature |
| MPI | Morphometric Protection Index |
| srad | Solar radiation (kJ $m^{-2}$ $day^{-1}$) |

Table S.2: Summary of the overlap coefficients between the eight duiker species. Upper triangle: posterior mean. Lower triangle: 90% posterior quantile.

| | *C. dorsalis* | *C. leucogaster* | *C. nigrifrons* | *C. rufilatus* | *C. silvicultor* | *C. weynsi* | *P. monticola* | *S. grimmia* |
|---|---|---|---|---|---|---|---|---|
| | | | | **$\bar{\Delta}_T$ : Overlap coefficient in time** | | | | |
| *C. dorsalis* | 0 | 0.285 | 0.338 | 0.235 | 0.752 | 0.176 | 0.202 | 0.571 |
| *C. leucogaster* | 0.196-0.383 | 0 | 0.482 | 0.632 | 0.384 | 0.632 | 0.595 | 0.524 |
| *C. nigrifrons* | 0.218-0.465 | 0.351-0.613 | 0 | 0.524 | 0.406 | 0.515 | 0.483 | 0.495 |
| *C. rufilatus* | 0.189-0.282 | 0.538-0.724 | 0.402-0.644 | 0 | 0.350 | 0.774 | 0.703 | 0.566 |
| *C. silvicultor* | 0.692-0.809 | 0.293-0.481 | 0.294-0.521 | 0.320-0.380 | 0 | 0.261 | 0.333 | 0.717 |
| *C. weynsi* | 0.137-0.220 | 0.539-0.721 | 0.400-0.628 | 0.737-0.810 | 0.235-0.289 | 0 | 0.747 | 0.427 |
| *P. monticola* | 0.157-0.250 | 0.511-0.681 | 0.370-0.597 | 0.678-0.728 | 0.305-0.362 | 0.722-0.772 | 0 | 0.53 |
| *S. grimmia* | 0.518-0.625 | 0.433-0.614 | 0.384-0.605 | 0.533-0.600 | 0.683-0.751 | 0.396-0.460 | 0.504-0.556 | 0 |
| | | | | **$\bar{\Delta}_S$ : Overlap coefficient in space** | | | | |
| *C. dorsalis* | 0 | 0.164 | 0.137 | 0.048 | 0.28 | 0.365 | 0.233 | 0.016 |
| *C. leucogaster* | 0.079-0.265 | 0 | 0.044 | 0.013 | 0.103 | 0.158 | 0.081 | 0.006 |
| *C. nigrifrons* | 0.049-0.250 | 0.008-0.112 | 0 | 0.072 | 0.158 | 0.171 | 0.194 | 0.045 |
| *C. rufilatus* | 0.028-0.074 | 0.004-0.030 | 0.028-0.133 | 0 | 0.308 | 0.065 | 0.131 | 0.307 |
| *C. silvicultor* | 0.210-0.352 | 0.053-0.165 | 0.075-0.252 | 0.265-0.351 | 0 | 0.35 | 0.332 | 0.202 |
| *C. weynsi* | 0.265-0.462 | 0.088-0.243 | 0.084-0.276 | 0.048-0.085 | 0.299-0.401 | 0 | 0.273 | 0.028 |
| *P. monticola* | 0.171-0.297 | 0.041-0.133 | 0.101-0.290 | 0.110-0.154 | 0.291-0.373 | 0.230-0.316 | 0 | 0.099 |
| *S. grimmia* | 0.007-0.030 | 0.001-0.017 | 0.011-0.105 | 0.228-0.383 | 0.155-0.249 | 0.017-0.042 | 0.079-0.122 | 0 |
| | | | | **$\bar{\Delta}_{ST}$ : Overlap coefficient in space and time** | | | | |
| *C. dorsalis* | 0 | 0.082 | 0.074 | 0.023 | 0.257 | 0.111 | 0.089 | 0.013 |
| *C. leucogaster* | 0.040-0.133 | 0 | 0.032 | 0.011 | 0.065 | 0.132 | 0.066 | 0.004 |
| *C. nigrifrons* | 0.027-0.136 | 0.006-0.080 | 0 | 0.054 | 0.099 | 0.123 | 0.131 | 0.033 |
| *C. rufilatus* | 0.014-0.036 | 0.004-0.026 | 0.021-0.100 | 0 | 0.166 | 0.062 | 0.117 | 0.233 |
| *C. silvicultor* | 0.193-0.322 | 0.034-0.105 | 0.047-0.159 | 0.144-0.190 | 0 | 0.147 | 0.168 | 0.185 |
| *C. weynsi* | 0.081-0.144 | 0.074-0.202 | 0.061-0.198 | 0.046-0.082 | 0.125-0.169 | 0 | 0.247 | 0.021 |
| *P. monticola* | 0.065-0.115 | 0.033-0.107 | 0.068-0.198 | 0.098-0.136 | 0.148-0.189 | 0.209-0.285 | 0 | 0.074 |
| *S. grimmia* | 0.006-0.025 | 0.001-0.013 | 0.008-0.077 | 0.174-0.289 | 0.142-0.228 | 0.013-0.032 | 0.059-0.09 | 0 |
